# One Health assessment of poultry and cattle farms as reservoirs for ESBL-producing *Escherichia coli*

**DOI:** 10.1101/2024.09.22.614308

**Authors:** Hina Malik, Randhir Singh, Simranpreet Kaur, Neha Parmar, Anuj Tyagi, R. S. Aulakh, J.P.S. Gill

## Abstract

**BACKGROUND:** Unregulated use of antibiotics in animal food production leads to the emergence of multidrug resistant bacteria, notably extended-spectrum beta-lactamases producing *Escherichia coli* (ESBL *E. coli*). ESBLs severely limit treatment options and pose a significant threat to public health. Investigating the potential role of livestock and poultry farms in harboring ESBL *E. coli* is crucial, given the prevalence of ESBL *E. coli* in humans, animals, and the environment, following the One Health framework.

**OBJECTIVES:** The aim of this study was to assess and compare the potential contribution of cattle and poultry farms along with their respective human handlers to the emergence and spread of ESBL *E. coli* following One Health approach.

**METHODS:** Twenty cattle and twenty poultry farms were surveyed in Ludhiana, India to determine antibiotic usage, prevalent diseases in the farms. Animal faeces/poultry droppings, handler hand swabs, and stool samples were examined for the presence of ESBL *E. coli*. The isolates were examined for their susceptibility to 16 antibiotics and for the presence of selected antimicrobial resistance genes, including ESBL genes. Generalized linear mixed models were applied to investigate variations in the prevalence of ESBL *E. coli* associated with farm animal species, farm practices, and farm biosecurity measures.

**RESULTS:** Our findings revealed a higher prevalence of ESBL *E. coli* in poultry farms (53.75%) than in cattle farms (40.72%) and overall lower prevalence (12%) in human handlers. Of the total isolates, 86.52% were multidrug resistant, and 93.91% showed a high multiple antibiotic resistance (MAR) index (>0.2). Isolates from poultry farms exhibited a higher level of resistance with 97% of isolates exhibiting multidrug resistance, 100% isolates with MAR index of more than 0.2, and 90.70% carrying the *bla*_CTXM_ gene.

**CONCLUSION:** Our findings highlight cattle and poultry farms as potential reservoirs for multidrug-resistant and ESBL *E. coli*, contributing to the rising environmental burden of antibiotic resistance genes. This emphasizes the urgency of limiting antibiotic use in animal farming, especially in poultry production, to address the escalating problem of antibiotic resistance and protect both animal and human health.

## INTRODUCTION

The emergence of beta-lactam antibiotic resistance within Enterobacteriaceae family is a major concern in the medical and veterinary fields (Wieler, 2014). This resistance is primarily due to the emergence of extended-spectrum beta-lactamases (ESBLs), which are enzymatic entities capable of hydrolyzing various antibiotics, including broad-spectrum cephalosporins and penicillins, and are encoded by genes such as *bla*_TEM_, *bla*_SHV_, and *bla*_CTXM_. Specifically, ESBL producing *Escherichia coli* (ESBL *E. coli*), a common species among ESBL-producing Enterobacteriaceae, causes serious health concerns in hospitals and communities (Peirano & Pitout, 2019). Infection by these multidrug resistant strains makes treatment difficult, and they are more likely to share their resistant traits with other bacterial species (McInnes et al., 2020). Clinically relevant *E. coli* strains affect animal, poultry, and the environment in addition to humans (Bevan et al., 2017). The extensive use of antibiotics in animals has been associated with the rise of antimicrobial-resistant bacteria in humans. This underscores the critical need to understand the distribution of ESBL *E. coli* and their associated drug-resistance genes across both human and animal populations following One Health approach owing to its far-reaching implications for ecological balance and human welfare (Tseng et al., 2023). Enterobacteriaceae that produce ESBLs frequently co-produce other beta-lactamases, such as AmpC lactamases, Metallo-beta lactamases (MBL), and *Klebsiella pneumoniae* carbapenemases (KPC), which confer resistance to third-generation cephalosporins and carbapenems. Thus, it is essential to assess the prevalence and threat of not only ESBL-producing but also other beta-lactamase-producing Enterobacteriaceae in livestock and poultry, following the One Health framework.

India stands out as a significant antimicrobial resistance (AMR) hotspot, characterized by alarming rates of AMR emergence (Zhao et al., 2024). Despite the predominant focus on the injudicious use of antimicrobials for human and animal health in most reports (Tang et al., 2023), the role of animals in AMR remains largely overlooked in India, perhaps due to the associated complexity in gathering such data. However, given the increased potential of transmission of multidrug resistant *E. coli* strains between humans and animals in India (Kuralayanapalya et al., 2019) and the significant roles of cattle and poultry farming in Indian agriculture sector, addressing multidrug resistant *E. coli* following One Health approach is of paramount importance. Nevertheless, while ESBL *E. coli* in animals have been limitedly studied, comprehensive knowledge of other beta-lactamase-producing *E. coli* in cattle and poultry, and in their human handlers is lacking in India. The goal of this study was: (i) to identify prevalent diseases and commonly used antibiotics in studied cattle and poultry farms through questionnaire-based survey; (ii) to identify ESBL, AmpC, MBL, and KPC producers in *E. coli* isolates from these farms; (iii) to identify and compare the antimicrobial resistance profiles of isolates from cattle, poultry, and farm workers; and (iv) to investigate variations in the prevalence of ESBL *E. coli* associated with farm animal species, practices, and farm biosecurity measures. The comprehensive exploration of ESBL *E. coli* in cattle, poultry, and animal handlers will provide valuable insights into resistance dynamics from a One Health perspective. Such findings will aid in developing targeted strategies to combat the dissemination of multidrug-resistant bacteria, ensuring both animal and public health.

## MATERIALS AND METHODS

### Survey

Punjab is one of the leading states in livestock and poultry production in India, thus, was chosen for this study. In order to identify prevalent diseases and commonly used antibiotics in both cattle and poultry and assess the biosecurity measures taken at the studied farms, a concise questionnaire was prepared for the owners of the selected farms. Farm owners were asked short and clear questions including demographic characteristics of farm owners (i.e. sex, age, occupation, level of education), type of farm (i.e. large farm > 50 cattle/ 5000 poultry > small farm), production method (i.e. intensive/semi-intensive/extensive), prevalent diseases at that time, commonly used antibiotics, antibiotic resistance understanding, and other farm practices, such as method of drug administration and disposal etc. Farm premises were investigated for the presence, storage, and disposal of antibiotics. The farm’s biosecurity measures were examined for internal (i.e. disease management, sanitation, and health management) and external biosecurity (i.e. infrastructure, location of farm, feed and water supply, removal of manure and dead animals, visitor and staff entry, vermin and vector control) by asking dichotomous questions. Based on the responses to questionnaire, the biosecurity score of each farm was determined by giving equal weight to each biosecurity measure and scoring each measure as either 1 or 0 following Kouam et al., (2018) which was then converted to percentage.

### Sample collection

Samples were collected from humans, cattle, and poultry in seven tehsils of Ludhiana district of Punjab after taking ethical permission under the reference no: DMCH/R&D/2021/7, Dated: 20/01/2021. Fresh faecal samples from animals were randomly collected from the ground after defecation. Utmost care was taken to avoid environmental contamination; after removing top layer, middle part was collected after proper mixing. Human samples were voluntarily provided (i.e. stool) and/or collected (i.e. hand swabs) by participating farm workers after signing declaration of consent. A total of 488 samples were collected from human farm workers, dairy cattle, and broiler chickens. Specifically, from cattle farms (large = 11, small = 9), a total of 248 samples were collected, comprising stool (n =15), and hand swab samples (n=33) from the farm workers and cattle faecal samples (n = 200). From poultry farms (large = 16, small = 4), a total of 240 samples were collected comprising stool (n = 6) and hand swab samples (n= 34) from poultry farm workers and poultry droppings (n = 200). All the samples were collected in Eppendorf tubes and kept in Styrofoam isolation box with ice and immediately transferred to the Centre for One Health, Guru Angad Dev Veterinary and Animal Sciences University for processing.

### Isolation and confirmation of Extended Spectrum Beta Lactamase producing *E. coli* (ESBL *E. coli*)

All the faecal samples were diluted in buffered peptone water (10%) in 1:9 dilution and inoculated on HiChrome™ ESBL agar (HiMedia, India) supplemented with HiCrome™ ESBL Selective Supplement (FD 278). The plate was then incubated under aerobic conditions at 37°C for 24-48 hours. Presumptive ESBL *E. coli* isolates, identified by their purple colonies, were subsequently subcultured on eosine methylene blue (EMB) agar. The taxonomic identities of grown colonies at species level were confirmed by Matrix Assisted Laser Desorption Ionization-Time of Flight Mass Spectrometry (MALDI-TOF MS) using an MALDI Biotyper® sirius system as described previously (Strejcek et al., 2018).

A modified CLSI combined disk test was performed to confirm ESBL production among *E. coli* isolates (Poulou et al., 2014). In brief, test culture with 0.5 McFarland turbidity was inoculated on Mueller Hinton Agar (MHA) and antibiotic disks (cefotaxime (CTX), ceftazidime (CAZ), cefotaxime-clavulanic acid (CEC) and ceftazidime-clavulanic acid (CAC)) impregnated with phenylboronic acid (PBA) (400 µg) and 0.1 M EDTA (292 µg) were placed on the cultured lawn. After overnight incubation at 37°C, the zone of inhibition around disks was measured. The tested isolate was considered as an ESBL producer if the difference between the zones of inhibition of CEC and CTX, and/or CAC and CAZ, was greater than 5 mm.

### AmpC and carbapenemase (*Klebsiella pneumoniae* carbapenemase (KPC), Metallo Beta Lactamase (MBL), and KPC-MBL co-producers) production test

To detect AmpC production in *E. coli* isolates, lawn of *E. coli* ATCC 25922 was inoculated on MHA. AmpC disks with 20 µL of 50:50 solutions of 100X tris-EDTA and normal saline, impregnated with test culture were placed adjacent to cefoxitin disk (FOX; 30 mcg), to examine the hydrolysis of cefoxitin due to the production of AmpC. Indentation or flattening of the cefoxitin inhibitory zone considered positive for AmpC production (Black et al., 2005).

To detect carbapenemase production, sterile meropenem (MEM; 10 µg) antibiotic disks were added to 2 ml of test culture broth (0.5 McFarland turbid) and incubated for 2-4 hours at 37°C. The disk then placed onto MHA plate inoculated with the lawn of *E. coli* ATCC 25922 and incubated at 37°C for 18 hours. The reduction in the zone of inhibition of the meropenem disk indicated the production of carbapenemases by the test culture (van der Zwaluw et al., 2015).

All the confirmed carbapenemase producers were further tested to identify type of carbapenemase being produced. Test culture (0.5 Mc Farland turbid) was inoculated on MHA on which four meropenem disks were placed, of which one was untreated, the other three disks were impregnated to get the combinations of meropenem + PBA (400 µg) (Tsakris et al., 2009), meropenem + EDTA (292 µg) (Yong et al., 2002), and meropenem + PBA (400 µg) + EDTA (292 µg) (Tsakris et al., 2010). Upon 18 hours incubation at 37°C, the zones of inhibition were compared. The type of carbapenemase produced was interpreted according to recommendations from previous studies (Tsakris et al., 2009, 2010; Yong et al., 2002).

### Antibiotic susceptibility test (ABST)

All ESBL *E. coli* isolates were assessed for the antibiotic susceptibility using a panel of 16 antibiotics representing 8 distinct antibiotic classes. The selection and categorization of antibiotics was based on the World Health Organization’s classification; highest priority critically important antibiotics, high priority critically important antibiotics, and highly important antibiotics (World Health Organization, 2018). The Kirby Bauer disk diffusion susceptibility protocol was used for ABST (Hudzicki, 2009) using commercially available antibiotic disks (HiMedia) by following Clinical Laboratory Standards Institute, 2020 (CLSI) guidelines (CLSI, 2020). In brief, the test cultures with 0.5 McFarland turbidity, were used for susceptibility to Ceftriaxone (CTR; 30 mcg), Ciprofloxacin (CIP; 5 mcg), Levofloxacin (LE; 5 mcg), Amikacin (AK; 30 mcg), Gentamicin (GEN; 10 mcg), Tobramycin (TOB; 10 mcg), Ampicillin (AMP; 10 mcg), Amoxicillin-clavulanic acid (AMC; 20/10 mcg), Meropenem (MRP; 10 mcg), Chloramphenicol (C; 30 mcg), Cefazolin (CZ; 30 mcg), Co-trimoxazole (COT; 25 mcg), Trimethoprim (TR; 5 mcg), and Tetracycline (TET; 30 mcg). Multidrug resistant strains were detected as described by Centers for Disease Control and Prevention (Magiorakos et al., 2012) and Multiple Antibiotic Resistance (MAR) index was calculated following Krumperman formula (Krumperman, 1983).

### DNA isolation and detection of selected antimicrobial resistance genes

DNA extraction of the freshly grown *E. coli* isolates was performed using the heat treatment method (Dashti et al., 2009). The supernatant containing DNA was collected, and quality and quantity of the genomic DNA was determined using NanoDrop spectrophotometer. The DNA samples were then stored at −20°C until further use.

PCR was performed in order to confirm the presence of 15 antimicrobial resistance genes in genomic DNA using primers pairs as listed in Table 1. A 10 µl PCR reaction volume, comprising 2.5 µl PCR master mix, 0.3 µl of each forward and reverse primers, 2.5 µl DNA template, and nuclease free water to make up the final volume, was subjected for amplification of each gene following the temperature, cycles parameters as described earlier (Table 1). The amplified products were analyzed using 1.5% agarose gel with ethidium bromide (0.5 μg/ml) and visualized using the gel documentation system (Syngene, USA).

**Table 1:**
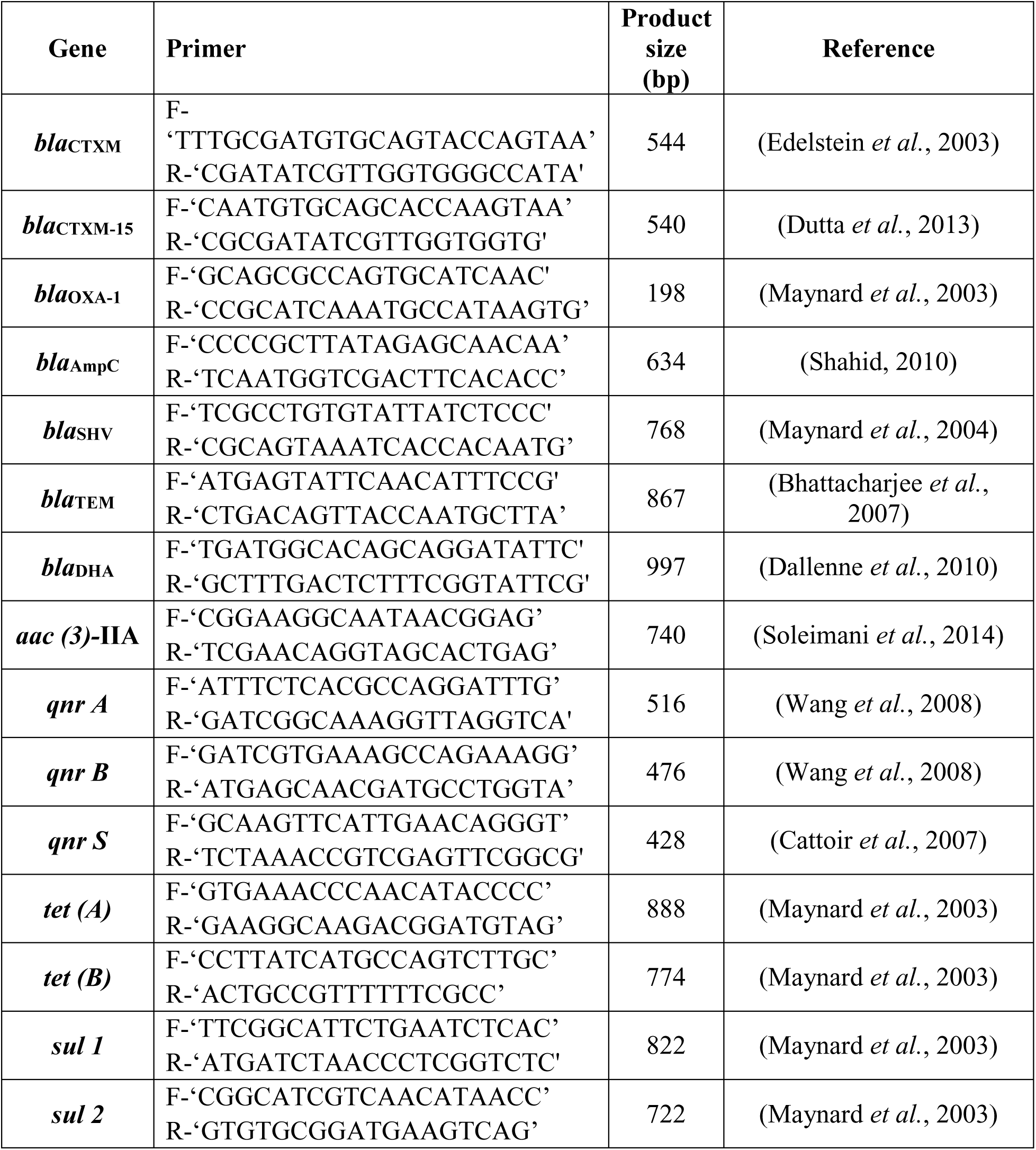
Primers used for the detection of the antimicrobial resistance genes in *E. coli*.

### Statistical Analysis

All statistical analyses were performed in the R environment (version 4.2.0). To determine inter-individual variability in the occurrence of ESBL *E. coli*; generalized linear mixed models (GLMMs) were built (*lme4*). To model ESBL *E. coli* occurrence, binomial family distribution with logit function (link = “logit”) was used. We tested the effect of farm species (Cattle; n=240, Poultry; n=240), farm type (Large; n=322, Small; n=158), and sample source (Faeces; n=400, Hand swabs; n=67, Human stool;n= 13) on calculated indices. In addition, we included the variables such as biosecurity score, owner education status, presence of antibiotics observed in the farm and treatment of animals by professional (Vet; n=300, Non-vet; n=180) as these factors could influence the calculated indices. To control the lack of independence between farm id and area (tehsil), we set farm id nested within area (tehsil) as random factor variables. Model selection was based on the information-theoretic (IT) approach using a second order Akaike’s information criterion corrected for small sample sizes (AIC_C_) as an information criterion and Akaike weights (ω) to determine model support. For all GLMMs, we report both conditional and marginal coefficients of determination of each model (R^2^_GLMM(c)_, which explains the variance of both the fixed and random factors, and R^2^_GLMM(m)_, which explains the variance of the fixed factors only), which we calculated as the variance explained by the best model and the ΔAIC_C_. The conditional parameter estimates (*β*), confidence intervals (95% CI) and odds ratio were calculated using model averaging with a cumulative AIC_C_ω of 95% (Burnham & Anderson, 2002). The sum of the AIC weights of each informative model in which a given predictor is included was used to calculate the relative importance of each predictor variable. Tukey’s General Linear Hypotheses Test (GLHT) was performed to compare average effects between groups when overall multivariable model significance was observed. Additionally, a stratified analysis approach, by separately analyzing cattle and poultry farms was employed, by building similar GLMM model for each species as above after removing farm species as variable. To test the correlation between phenotypic and genotypic antibiogram, a correlation matrix was plotted using the “corrplot” package (version 0.84; (Wei & Simko, 2017)). Principal correspondence analysis (PCA) was performed based on sensitivity zones of all tested antibiotics of the isolates (*FactoMineR*-version 4.2.1, (Lê et al., 2008). Following the PCA, Hierarchical Clustering on Principal Components (HCPC) was applied to cluster the ESBL *E. coli* isolates with similar phenotypic resistance profiles (*FactoMineR* - version 4.2.1, (Husson et al., 2010)). Similarly, to visualize genotypic antibiotic resistance pattern, multiple correspondence analysis was performed using 13 of the 15 tested genes since *qnr*A and *bla*_DHA_ had only one positive isolate for each gene (*FactoMineR*-version 4.2.1, (Husson et al., 2017)).

## RESULTS

We performed questionnaire-based survey mainly to identify prevalent diseases and commonly used antibiotics in farms as well as collected farm animals and their human handlers’ samples to evaluate the antibiotic resistance patterns of ESBL *E. coli* in the studied farms. We found that all cattle farms operated with intensive production systems, while all poultry farms used deep litter systems. The survey of both cattle and poultry farms revealed that the majority of farmers (i.e. owners, workers) in both types of farms were male, with all poultry farmers and 85% livestock farmers being male (Supplementary Table 1). Majority of the respondents were graduates, with 90% of poultry farmers and 50% of cattle farmers holding a graduate degree. The average age of farmer-owners was under 45, with 55% of them falling in this age group. For 75% of farmer-owners of the surveyed farms; animal farming was their major source of income. In cattle production, the most frequently reported diseases were mastitis, reproductive disorders, pyrexia of unknown origin, diarrhea, and respiratory disorder while in poultry, chronic respiratory disease, gumboro, and diarrhea were the most frequently reported diseases in the surveyed farms. Antibiotic use was prevalent in both cattle and poultry farming, with the majority of farmers (62%) seeking treatment from veterinarians and 38% turning to non-veterinarians. Interestingly, 78% of farms had at least one class of antibiotics in-use/stored for future use, and among these 55% had drugs from three or more different classes. Quinolones (70%) being the most frequently in-use/stored antibiotic class in both the farms, followed by aminoglycosides (19%), tetracycline (19%) and others (Supplementary Table 1). Interestingly, all these predominantly used antibiotics were found more in poultry farms than in cattle farms (quinolones; 65%-cattle, 75%-poultry, aminoglycosides; 35%-cattle, 60%-poultry, tetracycline; 35%-cattle, 60%-poultry).

### Occurrence of Extended Spectrum Beta Lactamase producing *E. coli* (ESBL *E. coli*)

A total of 230 ESBL *E. coli* were detected in 480 samples obtained from cattle and poultry farms all together. The frequency of occurrence of ESBL *E. coli* was higher in poultry farms than cattle farms (Table 2). In our run GLMM model for ESBL *E. coli* occurrence, a strong support for farm species, sample source, and biosecurity score was found in the best-fit model (ΔAIC_C_ = 67.74, R^2^_GLMM(m)_ = 0.241, R^2^_GLMM(c)_ = 0.296), using an information-theoretic (IT) approach. The relative importance of each predictor variable, as determined by the sum of AIC weights in averaged models, indicated that the biosecurity score (∑AIC_C_ω=1), farm species (∑AIC_C_ω=1) and sample source (∑AIC_C_ω=1) were equally important predictors, followed by the treatment prescription (i.e. treatment of animals by veterinarian/non-veterinarian) (∑AIC_C_ω=0.43) variable. However, farm type, owner education status, presence of antibiotics observed showed weak support in model outcome. The odds of farms with high biosecurity scores having ESBL *E. coli* were 0.93 (95% class interval: 0.90-0.96) times than those with lower biosecurity scores (Table 3, Supplementary Figure 1). Additionally, the odds of poultry farms having ESBL *E. coli* were 3.48 (95% class interval: 1.79-6.79) times higher than those of cattle farms (Figure 1A). Moreover, the presence of ESBL *E. coli* was significantly higher in animal faecal samples than in hand swab or human stool samples (Figure 1B). The odds of isolation of ESBL *E. coli* in hand swabs and human stool were 0.07 (95% CI: 0.03-0.17) and 0.25 (95% CI: 0.06-1.06), respectively.

**Figure 1:**
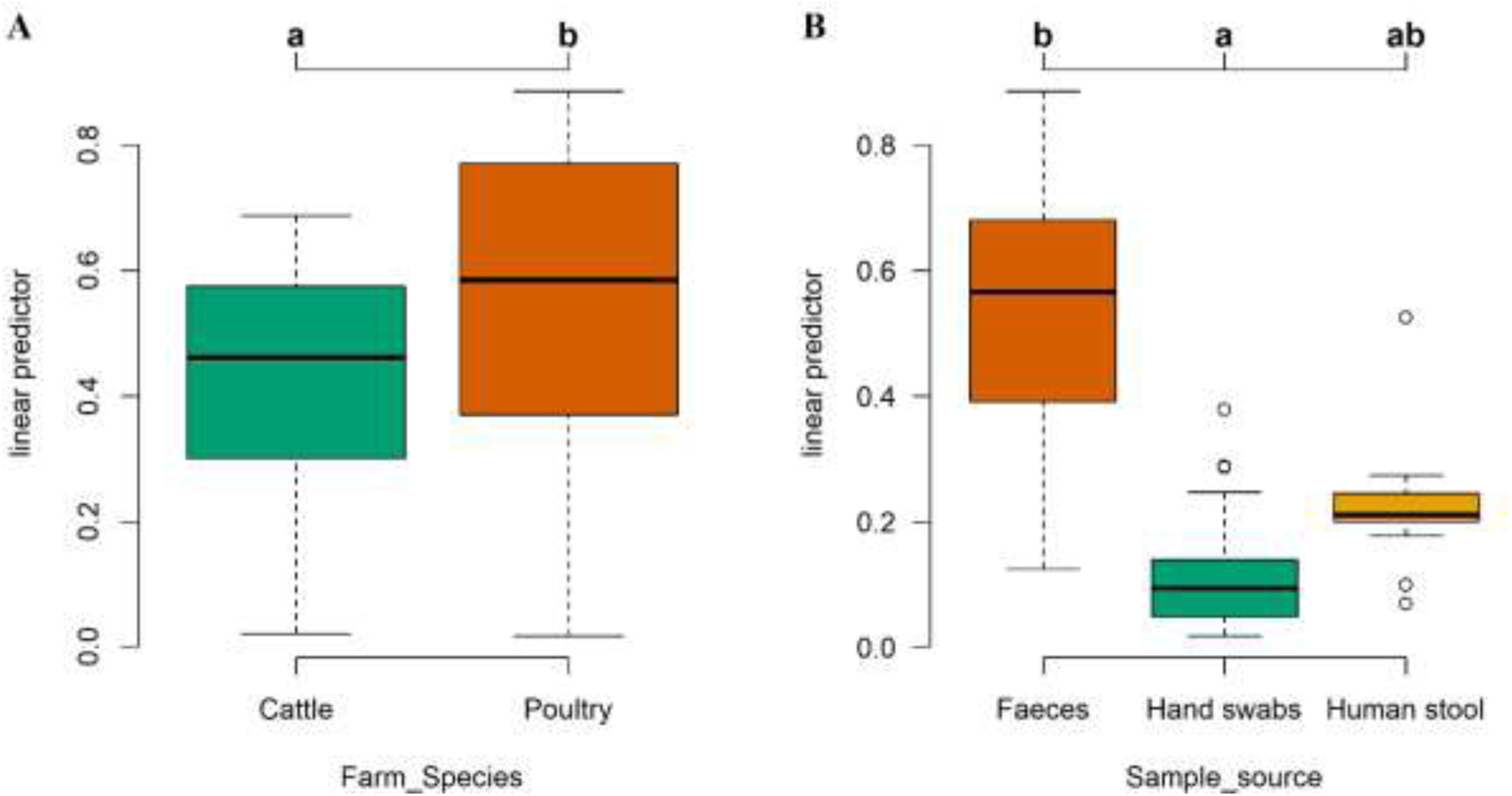
Comparison of ESBL *E. coli* occurrence between farm species (A) and sample source (B)

**Table 2:** Distribution of ESBL *E. coli* isolates in cattle and poultry farm samples.

| Source | Cattle | Poultry |
| --- | --- | --- |
| Animal faeces | 92/200 (46%) | 127/200 (63.5%) |
| Farm worker hand swabs | 6/33 (18.18%) | 2/34 (5.88%) |
| Farm worker stool | 3/15 (20%) | 0/6 (0%) |
| Grand Total | 101/248 (40.72%) | 129/240 (53.75%) |

**Table 3:** GLMM estimates and odds ratio of ESBL *E. coli* occurrence.

| Predictor variables | Estimate | Class Interval | p value | Odds Ratio | Class Interval |
| --- | --- | --- | --- | --- | --- |
| Biosecurity score* | -0.068 | (-0.098 to -0.038) | **<0.001** | **0.934** | (0.906 to 0.962) |
| Farm Species (Poultry)* | 1.249 | (0.582 to 1.916) | **<0.001** | **3.487** | (1.790 to 6.793) |
| Sample source (Hand swabs)* | -2.558 | (-3.399 to -1.716) | **<0.001** | **0.077** | (0.033 to 0.179) |
| Sample source (Human stool) | -1.377 | (-2.815 to 0.061) | 0.061 | 0.252 | (0.059 to 1.063) |
| Treatment (Vet) | -0.399 | (-1.001 to 0.203) | 0.194 | 0.671 | (0.367 to 1.225) |
| Owner graduation (Y) | -0.382 | (-1.020 to 0.256) | 0.240 | 0.682 | (0.360 to 1.291) |
| Antibiotics stored | -0.375 | (-1.135 to 0.384) | 0.333 | 0.687 | (0.321 to 1.468) |
| Farm type (Small) | -0.061 | (-0.821 to 0.698) | 0.874 | 0.940 | (0.439 to 2.011) |

### ESBL *E. coli* in Cattle farms

Within cattle farms, strong support for the effect of farm type (∑AIC_C_ω=1), sample source (∑AIC_C_ω=1), biosecurity score (∑AIC_C_ω=1), presence of antibiotics observed in the farm (∑AIC_C_ω=1) and treatment of animals by professional (∑AIC_C_ω=0.92), and moderate support for owner education status (∑AIC_C_ω=0.50) on the occurrence of ESBL *E. coli* was observed. Large cattle farms exhibited a higher occurrence (50.78%) of ESBL *E. coli* compared to small farms (30%) (p<0.01). Furthermore, farms where antibiotics were used or stored had a significantly higher likelihood (OR: 9.00; CI: 2.64-306.7, p<0.05) of harboring high levels of ESBL *E. coli*. Farms seeking treatment from veterinarians showed a lower probability (OR: 0.20; CI: 0.04-0.93, p<0.05) of containing high prevalence levels of ESBL *E. coli*.

### ESBL *E. coli* in Poultry farms

Within poultry farms, a strong support for an effect of sample source (∑AIC_C_ω=1), farm type (∑AIC_C_ω=0.69) on the occurrence of ESBL *E. coli* was observed similar to the omnibus model. Higher ESBL *E. coli* occurrence in samples from large farm compared to small farms, and in poultry samples compared to farm worker’s samples was noted. Intriguingly, the other investigated predictor variables, namely biosecurity score, antibiotic usage/storage, and treatment prescription, exhibited weak support on ESBL *E. coli* occurrence in poultry specific model.

### Antibiotic sensitivity pattern of the ESBL *E. coli* isolates

The antimicrobial resistance profile of ESBL *E. coli* isolates from both cattle and poultry farm samples was analyzed. Notably, a significant prevalence of resistance was observed across multiple antibiotic classes, with particularly high resistance to 3^rd^ generation cephalosporin, quinolones, and aminoglycosides (Supplementary Figure 2A, Supplementary Table 2). In cattle, resistance to beta-lactam, tetracycline and amikacin was prominent, while poultry samples exhibited high resistance to cephalosporins and quinolones. Importantly, 86.52% of the isolates were multidrug-resistant (MDR), with a higher proportion in poultry farms (96.90%) compared to cattle farms (73.27%). The majority of isolates (64.51%) showed resistance to at least four antibiotic classes (Supplementary Table 2). Among cattle farm isolates, multidrug resistance was observed in 72.83% of faecal isolates, 100% of human stool isolates, and 66.67% of hand swabs. Among poultry farm isolates, multidrug resistance was observed in 96.85% of droppings and 100% of hand swabs. Overall, 36 distinct MDR profiles were observed, with the most common profile (13%) being resistance to beta-lactams, quinolones, aminoglycosides, and tetracycline.

MAR index of ESBL *E. coli* isolates.

A significant percentage of isolates exhibited high multiple antibiotic resistance (MAR) values, with 93.91% (216/230) of the isolates showing a MAR index value >0.2. Particularly, 100% of poultry isolates and 86.14% (87/101) of cattle isolates showed MAR values greater than 0.2. Notably, human stool isolates had the highest MAR indices, followed by animal and human hand swab samples. Supplementary table 3 presents a summary of the MAR indices observed in the study, along with the corresponding number of isolates.

### Beta lactamase co-producers

Beta-lactamase enzymes other than ESBL were also investigated in 230 ESBL *E. coli* isolates, which revealed that 90, 1, and 2 isolates also produced AmpC, KPC, and KPC-MBL enzymes, respectively (Figure 2). Among the ESBL-AmpC co-producers, 43.41% (56/129) of isolates were from poultry, and 33.66% (34/101) were from cattle farms. Only one isolate from a large poultry farm was identified as an ESBL-KPC producer. Two ESBL-MBL-KPC co-producing *E. coli* were identified from poultry droppings.

**Figure 2:**
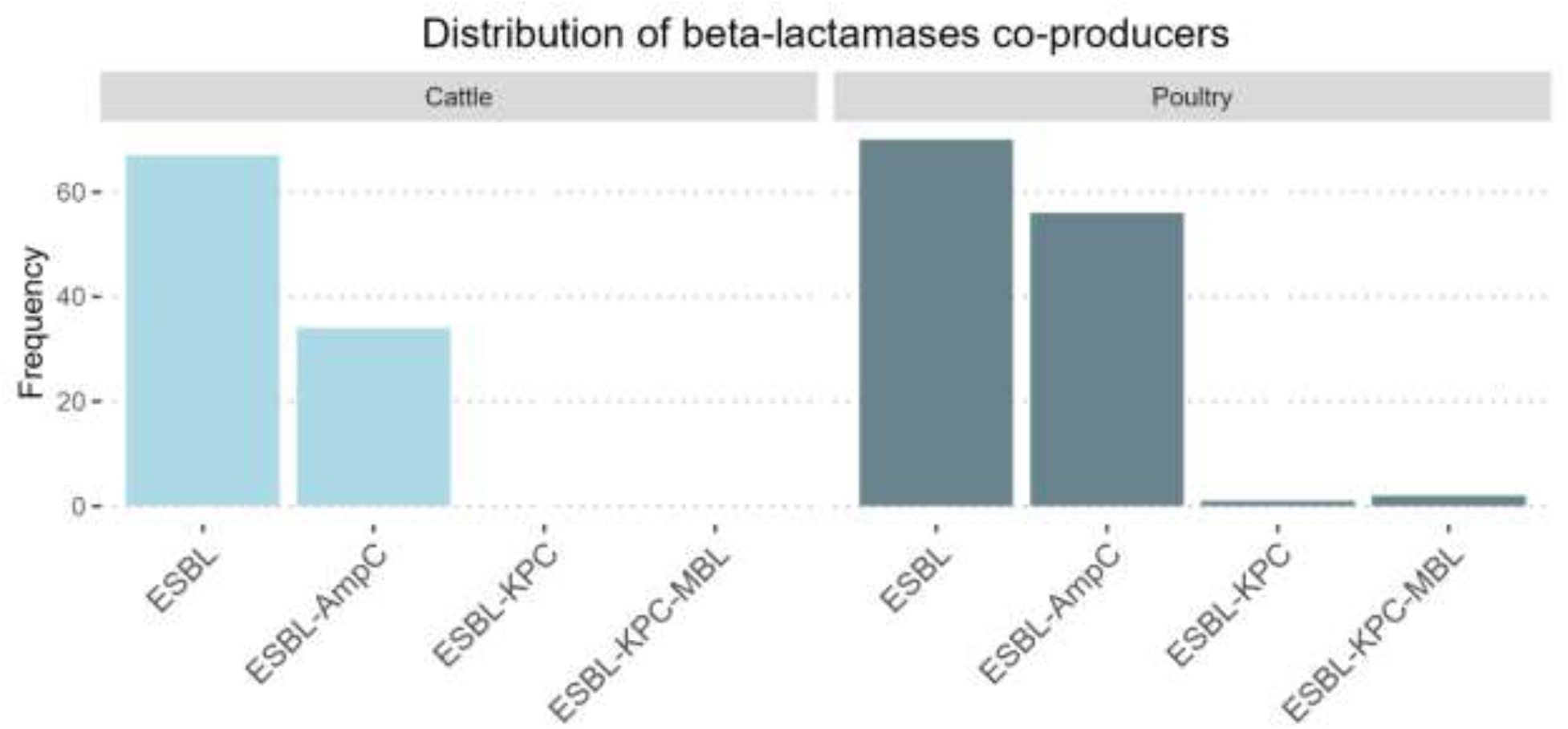
Frequency of *E. coli* isolates producing different combinations of beta in Cattle and Poultry

### Antimicrobial resistance genes in *E. coli* isolates

All the 230 ESBL *E. coli* isolates were examined for the presence of 15 antimicrobial resistance genes, which included beta-lactam genes (*bla*_OXA_, *bla*_CTXM_, *bla*_CTXM15_, *bla*_SHV_, *bla*_TEM_, *bla*_DHA_, and *bla*_AmpC_), an aminoglycoside gene (*aac*(3)-IIA), tetracycline genes (*tet*(A) and tet(B)), quinolone genes (*qnr*A, *qnr*B, and *qnr*S), and sulphonamide genes (*sul*1 and *sul*2) (Supplementary Figure 2B, Supplementary Table 4). The most abundant gene found in the ESBL *E. coli* isolates was *bla*_AmpC_, present in 83.48% (192) of the isolates, followed by *bla*_CTXM_ (80.87%), *bla*_TEM_ (57.39%), *qnr*S (37.83%), and *sul*2 (27.83%). Other beta-lactam genes such as *bla*_OXA_, *bla*_CTXM15_, *bla*_SHV_, and *bla*_DHA_ were found in 11.30%, 16.09%, 2.17%, and 0.43% of the isolates, respectively. In general, it was observed that the poultry isolates had a higher occurrence of antimicrobial resistance genes compared to cattle isolates, however there were gene specific differences. For example, the carriage of *bla*_SHV_ was higher in cattle isolates compared to poultry isolates, while the carriage of *bla*_CTXM_ was higher in poultry isolates.

### Association between antimicrobial resistance phenotype and resistance genes

Phenotypic antibiotic resistance was shown to have a significant positive correlation (p ≤0.05) with the presence of antimicrobial resistance genes, as depicted in Figure 3. In particular, a strong positive correlation was observed between the presence of *bla*_CTXM_ gene and resistance to ceftriaxone (r = 0.9), ampicillin (r = 0.9), and cefotaxime (r = 0.7); a moderate positive correlation with resistance to ceftazidime (r = 0.4); and a moderate negative correlation with the carriage of *bla*_SHV_ gene (r = −0.5). A moderate correlation between the presence of *sul*2 gene and sulphonamide resistance (r = 0.4) was observed, while a moderately positive correlation was found between the presence of *tet*A gene and *bla*_CTXM15_ (r = 0.6). A positive correlation between the resistance patterns of antibiotic classes is evident which supports the observation of multidrug resistance bacterial strains. In our study, we observed positive correlations in resistance patterns between the following antibiotics: betalactams with quinolones, tetracycline, and chloramphenicol; and between sulphonamides with tetracycline, and meropenem. Hierarchical clustering on PCA (HCPC) based on phenotypic resistance pattern (i.e. zone of inhibition) revealed three distinct clusters. (Figure 4A). The first cluster (n=38; P=29, C=5, CH=3, PH=1) was dominated by poultry isolates, which were phenotypically highly resistant to cephalosporin, quinolones, and aminoglycosides. The second and largest cluster (n=164; P=98, C=61, CH=4, PH=1) comprised most of the cattle and poultry isolates which were resistant to cephalosporin, quinolones, and tetracycline, but exhibited lower resistance to aminoglycosides. The third cluster was a separate group limited to cattle and cattle handlers’ isolates (n=28; P=0, C=26, CH=2, PH=0), which exhibited lesser resistance to cephalosporins, quinolones, and aminoglycosides. The clustering results suggest a differentiation in phenotypic resistance profiles between cattle and poultry isolates. Observed separation among clusters of bacterial isolates is due to broad differences in their antibiotic resistance patterns.

**Figure 3:**
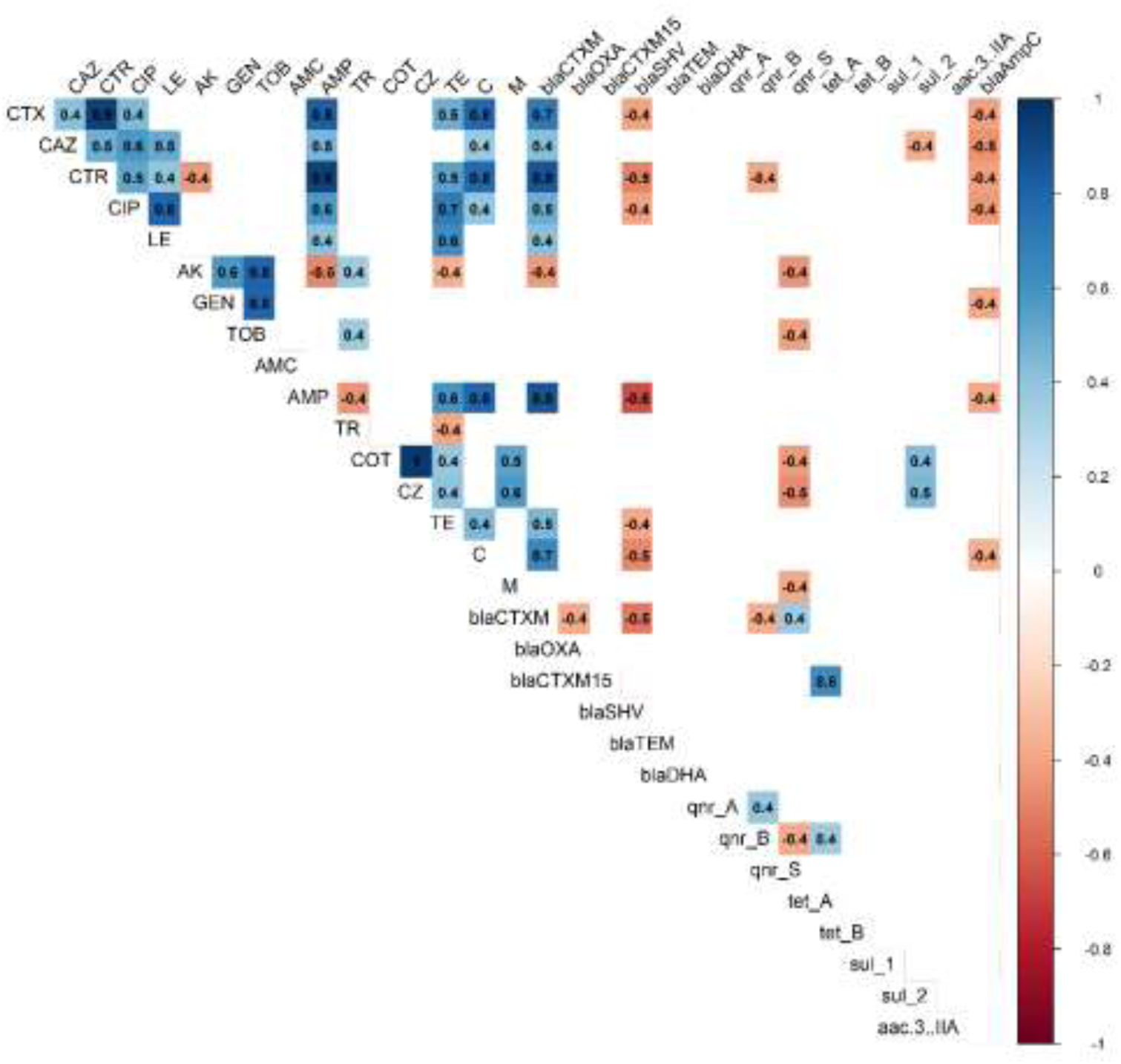
Correlation matrix showing significant correlation among phenotypic and genotypic resistance profiles along with r values

**Figure 4:**
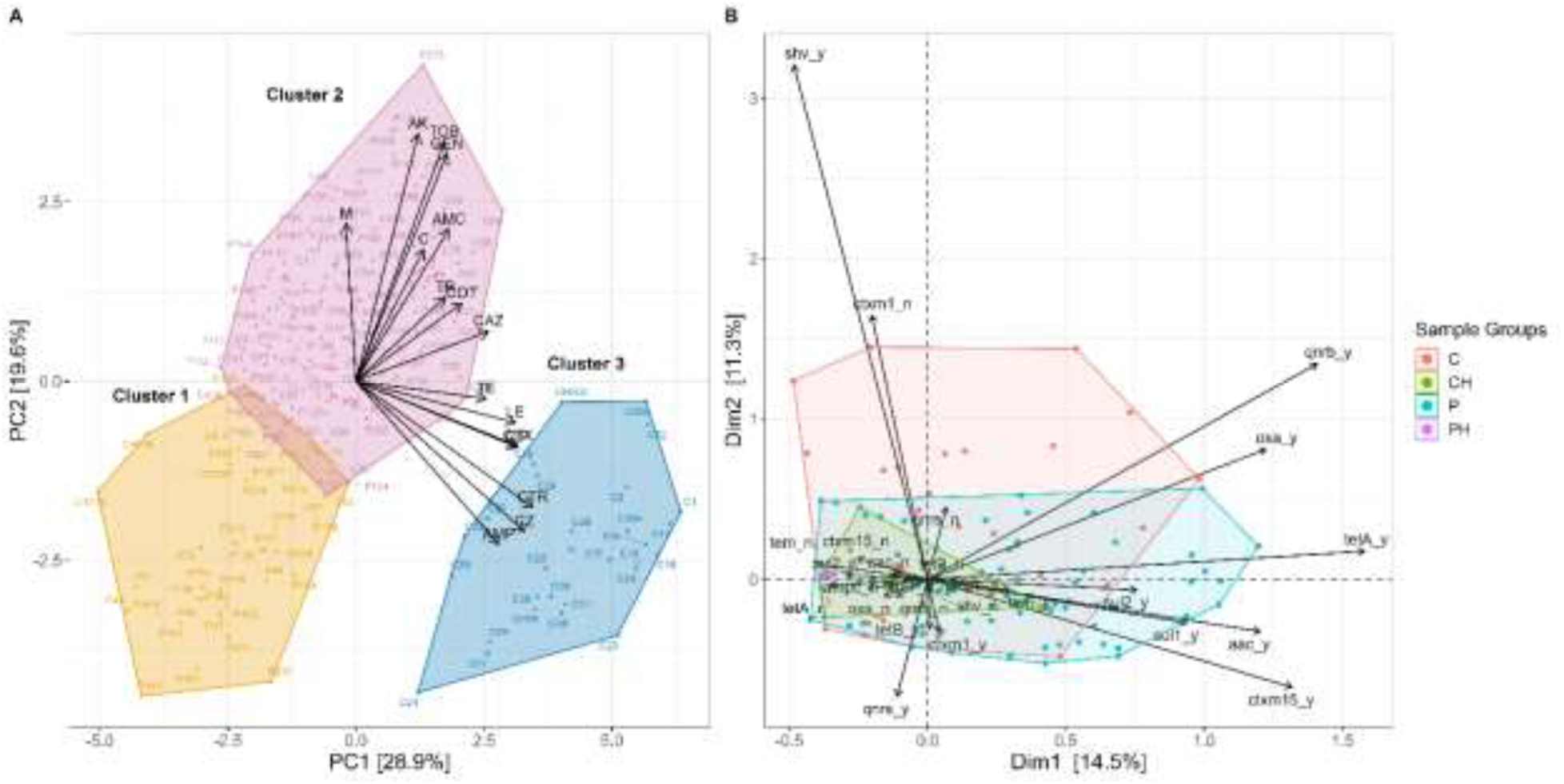
Hierarchical Clustering on Principal Component Analysis (HCPC) (panel A) and Multiple Correspondence Analysis (MCA) (panel B) of ESBL *E. coli* isolates with source labeling (Cattle [C], Cattle Handlers [CH], Poultry [P], Poultry Handlers [PH])

Similarly, MCA based on tested antibiotic resistance genes (i.e. AMR gene profile) revealed overlapping clusters showing differential spread of ESBL *E. coli* isolates mostly corresponding to their source of isolation. Observed clustering is majorly owing to higher prevalence of AMR genes such as *tet*(A), *bla*_CTXM15_, *qnr*B, *bla*_OXA_ and *sul*2, in poultry isolates, whereas higher prevalence of *bla*_SHV_ gene and absence of *bla*_CTXM_ gene in cattle isolates. For example, Dimension 1 was strongly influenced by the presence of *tet*A and *bla*_CTXM15_, while the absence of *bla*_CTXM_ and presence of *bla*_SHV_ had the greatest influence on dimension 2 (Figure 4B). The negative correlation between *bla*_SHV_ and *bla*_CTXM_ was also became apparent. Restricted spread of both poultry and cattle handler’s isolates indicates limited variability in their resistance pattern. Overall, we found high correlation between phenotype and genotype resistance pattern for isolates, which result in similarity in their phenotypic and genotypic clustering mostly reflecting their source of isolation.

## DISCUSSION

ESBL *E. coli* has gained importance in the recent past owing to their resistance to 3^rd^ generation cephalosporins leading to drug resistant infections in humans. We addressed ESBL *E. coli* prevalence, their phenotypic antimicrobial resistance pattern and antimicrobial resistance genes in different farm animals and their human handlers, following One-health framework in Punjab region of India. The findings of our survey demonstrate the extensive use of antibiotics in cattle and poultry farming, with multiple classes of antibiotics not only in use but also being stored for potential use in the future, indicating the possibility of the further development of antibiotic resistance. Restricting the spread of major prevalent diseases in the region, as noted in the survey, through good hygiene practices could not only limit the use of antibiotics but also lessen the financial burden on farmers, especially when animal farming is their sole occupation. Although the majority of farmers were educated, many still sought advice from non-veterinarians for animal treatment, which could have serious implications on animal health. Our study revealed that all the studied cattle and poultry farms in the region were ESBL *E. coli* positive and thus could serve as reservoirs of ESBL producing *E. coli* with the prevalence of 40% and 53.75%, respectively.

Several individual studies documented the occurrence of ESBL *E. coli* in either cattle or poultry production farms in India. Cattle farms have shown a lower prevalence of ESBL *E. coli*, ranging from 4.35% to 28% (Borah et al., 2014; Das et al., 2020), while the prevalence in poultry has been observed higher, from 6% to 87% (Brower et al., 2017; Chowdhury et al., 2022; Kar et al., 2015; Shrivastav et al., 2016). In comparison, lower prevalence rates of ESBL *E. coli* have been observed in cattle in America and European countries (Gelalcha et al., 2022; Li et al., 2018; Schmid et al., 2013; Weber et al., 2021). While a generally higher prevalence of ESBL *E. coli* in poultry compared to cattle is reported, significant variation in ESBL *E. coli* prevalence has been noted across different countries (De Koster et al., 2021; Liu et al., 2022; Mahamat et al., 2021). Raising poultry involves higher use of antibiotics than cattle production. For example, as per an estimate raising a kg of chicken annually requires more than thrice (148 mg/kg) the amount of antibiotics required to raise the same amount of cattle (45 mg/kg) produced annually (Van Boeckel et al., 2015). In our study also, we found that a higher percentage of poultry farms (85%) were using antibiotics for disease prevention or growth promotion compared to the cattle farms. Similarly, in a previous study from this region, 67% of poultry farmers admitted to using antibiotics as growth promoters (Brower et al., 2017). The study performed by Brower et al. (2017) in 2014 from Punjab region recorded 42% prevalence of ESBL *E. coli* in nine layer farms using phenotypic detection methods. After 8 years (2022) of this study, covering almost double number of farms (n=20), we found higher (53%) ESBL *E. coli* in poultry in the Punjab region of India associated with increased use of antibiotics (85%), emphasizing the seriousness of AMR problem in animal production sector in India.

In cattle farms, our data indicated that larger farm size was associated with a higher prevalence of ESBL *E. coli*, similar to what was observed in poultry farming, suggesting large intensive farming are at higher risk for ESBL *E. coli* compared to small scale farming as observed earlier by Gāliņa et al. (2021). This may be attributed to the greater density of animals and associated higher level of antibiotic use to control the diseases as noted by Wang et al. (2023). Additionally, cattle farms that stored antibiotics for future use and did not seek treatment from licensed veterinarians were more likely to harbor higher levels of ESBL *E. coli*. These issues, which have aggravated the AMR problem in India, have been discussed previously (Mutua et al., 2020). However, no appropriate measures have been taken so far to address the underlying factors, such as inadequate regulations and control over antibiotic sales (e.g. over-the-counter availability), limited access to veterinarians and the high cost of treatment. Overall, our study suggests that prudent use of antibiotics under professional supervision, along with implementing good biosecurity measures, may be crucial in reducing AMR burden in cattle farms.

Intriguingly, data from poultry farms suggest that the variables such as biosecurity score, antibiotic usage/storage and treatment prescription did not show a statistically significant impact on ESBL *E. coli* occurrence. The could be due to the fact that in our survey, 85% of the poultry farms were using antibiotics in disease prevention or in starter feed, supposedly for growth promotion. Such high use of antibiotics limits the ability of the statistical assessment of other factors being tested. Thus, the substantial prevalence of MDR ESBL *E. coli* in poultry droppings can be attributed to the intense selection pressure from the high use of antibiotics in poultry for disease prevention and growth promotion. ESBL *E. coli* isolates especially from poultry droppings, were found to co-produce other beta lactamase enzymes such as AmpC, KPC and MBL. Among these, MBL and KPC producing strains are of critical concern due to their extensive resistance profiles, rendering them completely resistant to all beta lactams and leaving severely limited treatment options. Additionally, resistance genes for these enzymes are plasmid-mediated, facilitating their spread and posing significant clinical and public health implications (Bush & Jacoby, 2010). Overall, given the extensive use of antibiotics, high prevalence of ESBL *E. coli*, and their broad resistance profiles observed in our study, it is evident that strategies to mitigate the spread of resistant bacteria in poultry farms must prioritize the judicious use of critically important antibiotics. This should be complemented by enhanced environmental management practices, regular monitoring, and increased awareness of farmers on responsible use of antibiotics.

In our study, we observed a high prevalence (93.91%) of MDR, especially among poultry isolates (100%). Earlier studies documented varying MDR; ranging between 30 - 89% in ESBL *E. coli* isolates from poultry (De Koster et al., 2021; Sonola et al., 2021; Tansawai et al., 2019). Compared to the present study, Waade et al. (2021) documented lower MDR in ESBL *E. coli* isolates (53%) from the cattle farms in Germany indicating lower use of antibiotics in those cattle farms. We noted positive correlation between the resistant phenotype of different antibiotic classes in ESBL *E. coli* isolates similar to other studies (Alwash & Al-Rafyai, 2019; Nossair et al., 2022; Sonola et al., 2021), indicating that resistance for critically important antibiotics could be influenced by the development of resistance to common antibiotics. Nearly all the ESBL *E. coli* isolates (99.13%) were resistant to at least one class of antibiotic, with the most commonly observed resistance to cephalosporin and tetracycline. This corroborates our survey findings where tetracycline and quinolones were the most commonly used preventive and therapeutic alternatives in the farms, along with 3^rd^ and 4^th^ generation cephalosporin, especially in cattle farms. The high prevalence of MDR in ESBL *E. coli* isolates might be attributed to factors such as high antimicrobial usage due to poor regulations and over-the-counter accessibility, as well as sharing of ARGs among microbial communities in the gut and the environment.

We observed three distinct clusters of phenotypic resistance among ESBL *E. coli* isolates using HCPC, distributed across both cattle and poultry. The first cluster was dominated by poultry isolates, the second cluster comprised most of the cattle and poultry isolates, and the third cluster was a separate group comprising of isolates from cattle and cattle handlers. This distinction likely reflects varying antibiotic usage practices in cattle and poultry farming, which contributed to different resistance profiles. Performed MCA elucidated the genotypic basis of these phenotypic patterns, specifically demonstrated, how specific ARGs influenced resistance profiles and contributed to segregation of cattle and poultry isolates into different overlapping groups. The correlation analysis between phenotypic antibiotic resistance and ARGs presence revealed significant positive correlations. The *bla*_CTXM_ gene showed a strong positive correlation with resistance to ceftriaxone, ampicillin, and cefotaxime, indicating its critical role in conferring resistance to these antibiotics. Interestingly, it was notable that most cattle isolates carrying *bla*_SHV_ were grouped together and remained sensitive to third generation cephalosporins, whereas *bla*_CTXM_ was more prevalent in poultry isolates, correlating with their resistance to these antibiotics, particularly cefotaxime. This finding is consistent with previous studies documenting carriage of ESBL genes such as *bla*_CTXM_, *bla*_SHV_ and *bla*_TEM,_ providing varied resistance to cephalosporin in cattle and poultry *E. coli* isolates (Ahmed et al., 2020; Cantón et al., 2012; Elmonir et al., 2021; Nossair et al., 2022; Tansawai et al., 2019). Additionally, a moderate positive correlation was observed between *tet*(A) gene presence and *bla*_CTXM15_, suggesting co-selection mechanisms.

Farm animal handlers working in close association with cattle and poultry are at risk of acquiring resistant strains of microorganisms. In the present study, we found 12% of samples from farm handlers were positive for ESBL *E. coli*. Although the prevalence of ESBL *E. coli* was higher in poultry than in cattle, interestingly the poultry farm workers had a lower (5%) prevalence than the dairy farm workers (18.75%). This could be due to the fact that dairy farm workers are more frequently involved in handling for feeding, milking and cleaning of dairy cattle than poultry, which put them at higher risk of acquiring ESBL *E. coli.* The hesitation of farm workers to share their stool samples for the study limited our in-depth investigation. Earlier studies that examined ESBL *E. coli* on dairy and poultry farms have also found human handlers positive for the ESBL *E. coli* (Dahms et al., 2015; Tansawai et al., 2019). For example, in a recent study, Nossair et al. (2022) found 20% of the stool samples were positive for ESBL *E. coli* from livestock and poultry farm workers. The prevalence of ESBL *E. coli* in farm workers might be attributed to a variety of factors, including, inadequate hygiene standards, and close contact between animals and people. The transfer of ESBL *E. coli* from animals to human handlers or vice-versa could be better addressed by knowing the specific ESBL *E. coli* strains and their abundance. Nevertheless, the consequences of ESBL *E. coli* carriage in healthy humans that too in farm workers could turn serious; for instance, it is estimated that nearly 250,000 people worldwide lose their life annually from resistant *E. coli* infections in hospitals (Murray et al., 2022). Resistance to meropenem was found in 9% of farmworker isolates, highlighting a serious threat. Similar patterns of resistance in ESBL *E. coli* have been documented in other studies (Nossair et al., 2022; Tansawai et al., 2019). The farm workers’ isolates in our study, also revealed *bla*_CTXM_ gene predominance which corroborates with the findings of a study from Europe (Dahms et al., 2015), while even higher carriage of the gene was observed in farmworkers of Egypt and Thailand (Nossair et al., 2022; Tansawai et al., 2019). The *bla*_CTXM_ gene governs the resistance to 3^rd^ generation cephalosporin which are prioritized by WHO as highest priority antibiotics for human use. Increased resistance to these antibiotics not only limits the infection control strategies options but also makes treatment costlier. The increased prevalence of *bla*_CTXM_ is a cause for further concern as these genes are plasmid-mediated and tend to disseminate among bacterial communities.

Overall our findings highlight the complex interplay between phenotypic and genotypic resistance in ESBL *E. coli* isolates in cattle and poultry, emphasizing the importance of tailored strategies to address antimicrobial resistance in these systems. In cattle farms, prioritizing biosecurity measures and ensuring antibiotic treatment on veterinarian prescription is crucial for reducing ESBL prevalence. While in poultry farms, stringent regulations on antibiotic use and improved waste management are essential to mitigate the selection pressure exerted by heavy antibiotic use aimed at preventing AMR burden.

## CONCLUSION

This study emphasizes that cattle and poultry farms play as reservoirs for multidrug-resistant and extended-spectrum beta-lactamase *E. coli*. Our data show a significantly greater occurrence of ESBL *E. coli* in poultry farms than in cattle farms, emphasizing poultry production as a significant contributor to the environmental burden of antibiotic resistance bacterial strains as well as antimicrobial resistance genes. The alarming levels of multidrug resistance and high multiple antibiotic resistance indices observed in cattle and poultry isolates, primarily driven by the widespread use of antibiotics, raise pressing concerns regarding the limited availability of effective treatment. This study highlights the crucial insights for implementing strategies, such as stringent regulations on antibiotic usage, enhanced biosecurity measures and public awareness, in order to address this escalating problem and protect both animal well-being and human health.

## Ethics Statement

The study was reviewed and approved by the Ethics Committee of Dayanand Medical College and Hospital, under the reference no: DMCH/R&D/2021/7, Dated: 20/01/2021.

## Author Contributions

HM and RS conceived the study. HM carried out laboratory experiments, statistical analyses, and wrote the manuscript. NP contributed in the sample collection, and RS edited the manuscript. All authors contributed to the article and approved the submitted version.

## Acknowledgments

We are grateful to Wasimuddin, Senior Scientist, Norwegian Veterinary Institute, for his scientific advice and all farmers for their constant help and support during our field work.

## Supplementary Material

The Supplementary Material for this article can be found with the article.

