## Supplementary file for "One Health assessment of poultry and cattle farms as reservoirs for ESBL-producing *Escherichia coli*"

J.P.S. Gill^1^

^1^Centre for One Health, Guru Angad Dev Veterinary and Animal Sciences University, Ludhiana,

Punjab, India

^2^College of Fisheries Sciences, Guru Angad Dev Veterinary and Animal Sciences University,

Ludhiana, Punjab, India

***Corresponding Author:**

Randhir Singh

Professor, Centre for One Health, Guru Angad Dev Veterinary and Animal Sciences University,

Ludhiana, Punjab, India

**Figures**

- Supplementary Figure 1: Relationship between ESBL *E. coli* occurrence and biosecurity score of the farms
- Supplementary Figure. 2: Comparison of antibiotic resistance patterns (A) and AMR gene distribution (B) in ESBL-producing *E. Coli* from cattle, poultry, and farm handlers

**Tables**

- Supplementary Table 1: Farm owner demographics, antibiotic practices, and common disease patterns: survey overview
- Supplementary Table 2: Antimicrobial resistance profile of ESBL *E. coli* isolates from cattle and poultry
- Supplementary table 3: Multi antibiotic resistance (MAR) index of ESBL *E. coli* isolates from cattle and poultry
- Supplementary table 4: Presence of antibiotic resistance genes (ARGs) in ESBL *E. coli* isolates from cattle and poultry

**Figures**

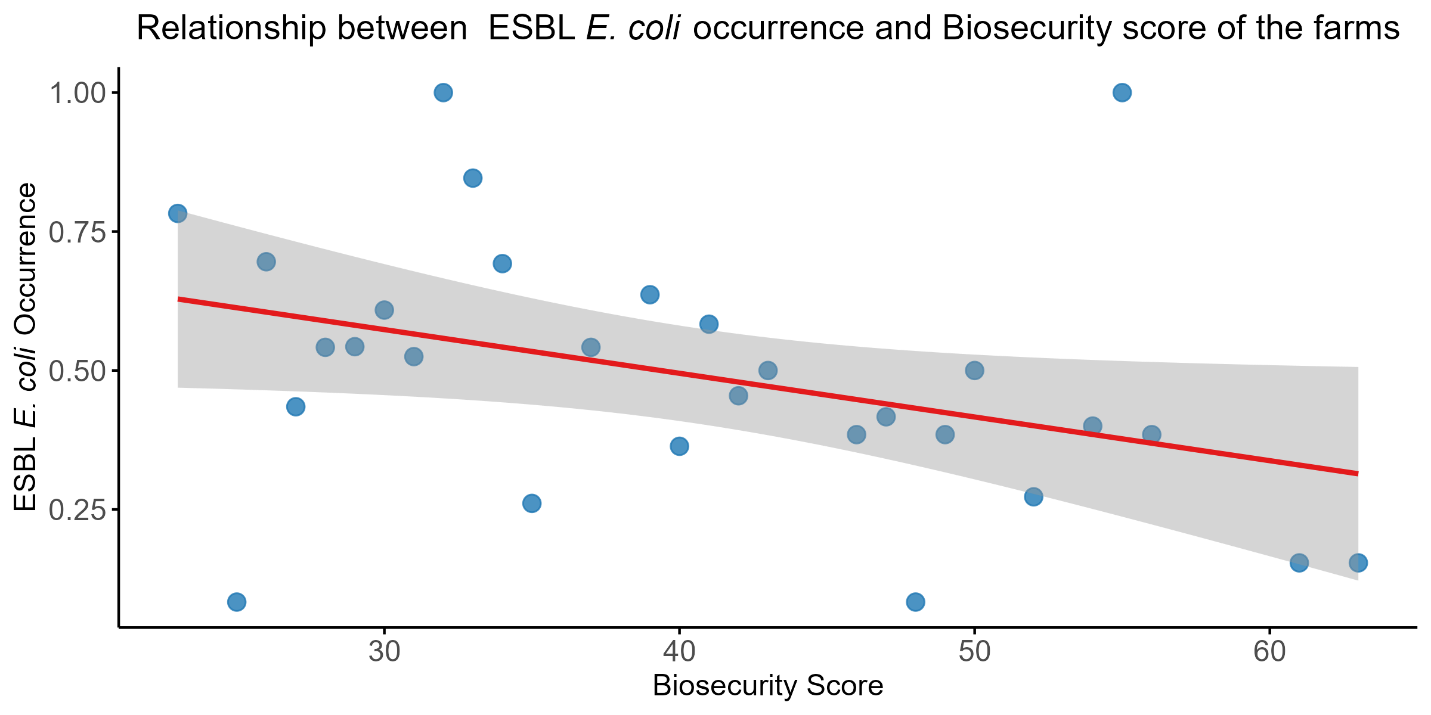

Supplementary Figure 1: Relationship between ESBL *E. coli* occurrence and biosecurity score of the farms

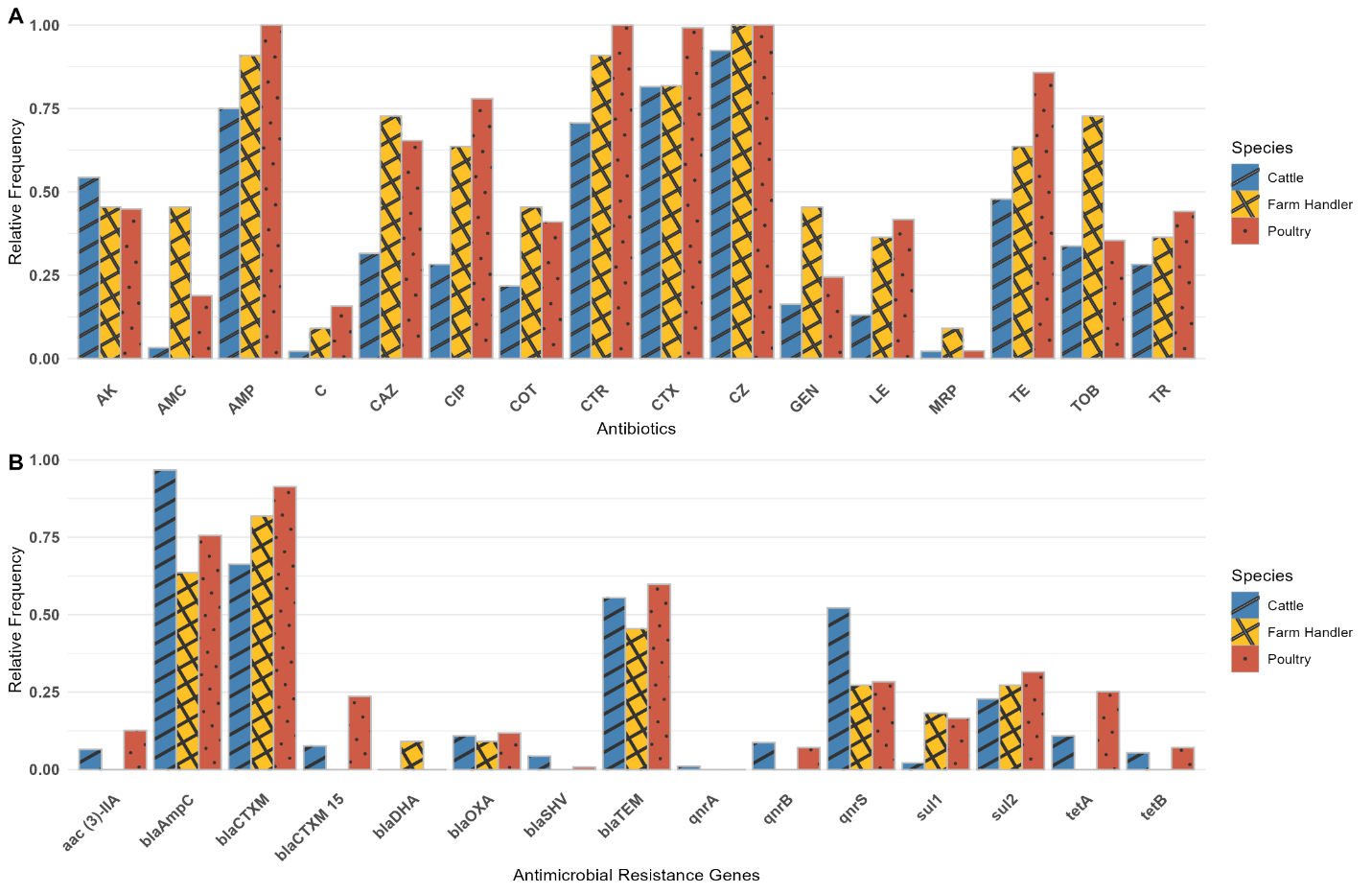

Supplementary Figure. 2: Comparison of antibiotic resistance patterns (A) and AMR gene distribution (B) in ESBL-producing *E. Coli* from cattle, poultry, and farm handlers

**Tables**

Supplementary Table 1: Farm owner demographics, antibiotic practices, and common disease patterns: survey overview

| **Variables** | **Response** | **Studied farms** | | |
| --- | --- | --- | --- | --- |
|  |  | **Cattle (n=20)** | **Poultry (n=20)** | **Total (n=40)** |
| Gender | Male | 17 (85.00%) | 20 (100.00%) | 36 (90.00%) |
|  | Female | 3 (15.00%) | 0 (0%) | 4 (10.00%) |
| Age | 30-45 | 9 (45.00%) | 13 (65.00%) | 22 (55.00%) |
|  | 46-60 | 5 (25.00%) | 7 (35.00%) | 12 (30.00%) |
|  | >60 | 6 (30.00%) | 0 (0.00%) | 6 (15.00%) |
| Education | Graduate | 10 (50.00%) | 18 (90.00%) | 28 (70.00%) |
|  | Undergraduate | 10 (50.00%) | 2 (10.00%) | 12 (30.00%) |
| Income share | Major | 14 (70.00%) | 16 (80.00%) | 30 (75.00%) |
|  | Medium to minor | 6 (30.00%) | 4 (20.00%) | 10 (25.00%) |
| Farming | Contract | 1 (5.00%) | 14 (70.00%) | 15 (37.50%) |
|  | Self-owned | 19 (95.00%) | 6 (30.00%) | 25 (62.50%) |
| Treatment by | Veterinarian | 10 (50.00%) | 15 (75.00%) | 25 (62.50%) |
|  | Non veterinarian | 10 (50%) | 5 (25.00%) | 15 (37.50%) |
| Antibiotics use/stored | Yes | 14 (70.00%) | 17 (85.00%) | 31 (77.50%) |
|  | No | 6 (30.00%) | 3 (15.00%) | 9 (22.50%) |
| Common diseases | Mastitis | 17 (85.00%) | NA | NA |
|  | Reproductive disorders | 16 (80.00%) | NA | NA |
|  | Pyrexia | 15 (75.00%) | NA | NA |
|  | Diarrhoea | 8 (40.00%) | 3 (15.00%) | 11 (27.50%) |
|  | Respiratory disorder | 6 (30.00%) | NA | NA |
|  | Chronic respiratory disease | NA | 18 (90.00%) | NA |
|  | Gumboro | NA | 7 (35.00%) | NA |
| Commonly stored antibiotics | Quinolones | 13 (65.00%) | 15 (75.00%) | 28 (70.00%) |
|  | Aminoglycosides | 7 (35.00%) | 12 (60.00%) | 19 (47.50%) |
|  | Penicillin | 8 (40.00%) | 0 (0.00%) | 8 (20.00%) |
|  | 3rd gen cephalosporin | 9 (45.00%) | 0 (0.00%) | 9 (22.50%) |
|  | Tetracycline | 7 (35.00%) | 12 (60.00%) | 19 (47.50%) |
|  | 2nd gen cephalosporin | 0 (0.00%) | 2 (10.00%) | 2 (5.00%) |
|  | Polymyxin | 1 (5.00%) | 0 (0.00%) | 1 (2.50%) |
|  | Sulphonamide | 2 (10.00%) | 0 (0.00%) | 2 (5.00%) |

Supplementary Table 2: Antimicrobial resistance profile of ESBL *E. coli* isolates from cattle and poultry

| **WHO categories** | | | **Highest priority Critically important Antibiotics** | | | | | **High priority Critically Important Antibiotics** | | | | | | **Highly important antibiotics** | | | | |
| --- | --- | --- | --- | --- | --- | --- | --- | --- | --- | --- | --- | --- | --- | --- | --- | --- | --- | --- |
| **Antibiotic classes** | | | **Cephalosporins (3rd gen)** | | | **Quinolones** | | **Aminoglycosides** | | | **Penicillin** | | **Carbapenem** | **Sulphonamides** | | **Tetracycline** | **Cephalosporin (1st gen)** | **Amphenicols** |
| **Farm** | **Source** | **Total isolates** | **Antibiotic disks (% resistance)** | | | | | | | | | | | | | | | |
|  |  |  | **CTX** | **CAZ** | **CTR** | **CIP** | **LE** | **AK** | **GEN** | **TOB** | **AMC** | **AMP** | **MRP** | **TR** | **COT** | **TE** | **CZ** | **C** |
| Cattle | Faeces | 92 | 75 (81.52) | 29 (31.52) | 65 (70.65) | 26 (28.26) | 12 (13.04) | 50 (54.35) | 15 (16.30) | 31 (33.70) | 3 (3.26) | 69 (75.00) | 2 (2.17) | 26 (28.26) | 20 (21.74) | 44 (47.83) | 85 (92.39) | 2 (2.17) |
|  | Hand swabs | 6 | 4 (66.67) | 4 (66.67) | 5 (83.33) | 2 (33.33) | 1 (16.67) | 2 (33.33) | 1 (16.67) | 3 (50.00) | 1 (16.67) | 5 (83.33) | 1 (16.67) | 2 (33.33) | 2 (33.33) | 3 (50.00) | 6 (100.00) | 0 (0.00) |
|  | Human stool | 3 | 3 (100.00) | 3 (100.00) | 3 (100.00) | 3 (100.00) | 2 (66.67) | 2 (66.67) | 2 (66.67) | 3 (100.00) | 2 (66.67) | 3 (100.00) | 0 (0.00) | 2 (66.67) | 3 (100.00) | 2 (66.67) | 3 (100.00) | 1 (33.33) |
|  | Total | 101 | 82 (81.19) | 36 (35.64) | 73 (72.28) | 31 (30.69) | 15 (14.85) | 54 (53.47) | 18 (17.82) | 37 (36.63) | 6 (5.94) | 77 (76.24) | 3 (2.97) | 30 (29.70) | 25 (24.75) | 49 (48.51) | 94 (93.07) | 3 (2.97) |
| Poultry | Faeces | 127 | 126 (99.21) | 83 (65.35) | 127 (100.00) | 99 (77.95) | 53 (41.73) | 57 (44.88) | 31 (24.41) | 45 (35.43) | 24 (18.90) | 127 (100.00) | 3 (2.36) | 56 (44.09) | 52 (40.94) | 109 (85.83) | 127 (100.00) | 20 (15.75) |
|  | Hand swabs | 2 | 2 (100.00) | 1 (50.00) | 2 (100.00) | 2 (100.00) | 1 (50.00) | 1 (50.00) | 2 (100.00) | 2 (100.00) | 2 (100.00) | 2 (100.00) | 0 (0.00) | 0 (0.00) | 0 (0.00) | 2 (100.00) | 2 (100.00) | 0 (0.00) |
|  | Human stool | 0 | 0 (0.00) | 0 (0.00) | 0 (0.00) | 0 (0.00) | 0 (0.00) | 0 (0.00) | 0 (0.00) | 0 (0.00) | 0 (0.00) | 0 (0.00) | 0 (0.00) | 0 (0.00) | 0 (0.00) | 0 (0.00) | 0 (0.00) | 0 (0.00) |
|  | Total | 129 | 128 (99.22) | 84 (65.12) | 129 (100.00) | 101 (78.29) | 54 (41.86) | 58 (44.96) | 33 (25.58) | 47 (36.43) | 26 (20.16) | 129 (100.00) | 3 (2.33) | 56 (43.41) | 52 (40.31) | 111 (86.05) | 129 (100.00) | 20 (15.50) |
| Grand Total | | 230 | 210 (91.30%) | 120 (52.17) | 202 (87.83) | 132 (57.39) | 69 (30.00) | 112 (48.70) | 51 (22.17) | 84 (36.52) | 32 (13.91) | 206 (89.57) | 6 (2.61) | 86 (37.39) | 77 (33.48) | 160 (69.57) | 223 (96.96) | 23 (10.00) |

Supplementary table 3: Multi antibiotic resistance (MAR) index of ESBL *E. coli* isolates from cattle and poultry

| Farm | **Source** | **MAR index** | | | | | | | | **Isolates with >0.2 MAR index** |
| --- | --- | --- | --- | --- | --- | --- | --- | --- | --- | --- |
|  |  | **0.2 - 0.29** | **0.3 - 0.39** | **0.4 - 0.49** | **0.5- 0.59** | **0.6 - 0.69** | **0.7 - 0.79** | **0.8 - 0.89** | **0.9- 0.99** |  |
| Cattle | Faeces (92) | 12 | 23 | 18 | 19 | 6 | 1 | 0 | - | 79 (85.87%) |
|  | Hand swabs (6) | - | 3 | - | 0 | 1 | 1 | 0 | - | 5 (83.33%) |
|  | Human stool (3) | - | 0 | - | 0 | 1 | 1 | 0 | 1 | 3 (100.00%) |
|  | Total (101) | 12 | 26 | 18 | 19 | 8 | 3 | 0 | 1 | 87 (86.14%) |
| Poultry | Faeces (127) | 4 | 18 | 12 | 39 | 35 | 9 | 9 | 1 | 127 (100.00%) |
|  | Hand swabs (2) | - | 0 | - | 1 | 0 | 1 | 0 | - | 2 (100.00%) |
|  | Total (129) | 4 | 18 | 12 | 40 | 35 | 10 | 9 | 1 | 129 (100.00%) |
| Grand Total (230) | | 16 | 44 | 30 | 59 | 43 | 13 | 9 | 2 | 216 (93.91%) |

Supplementary table 4: Presence of antibiotic resistance genes (ARGs) in ESBL *E. coli* isolates from cattle and poultry

| **Farm** | **Source** | **Total isolates** |  | **Antimicrobial resistance genes (%)** | | | | | | | | | | | | | |
| --- | --- | --- | --- | --- | --- | --- | --- | --- | --- | --- | --- | --- | --- | --- | --- | --- | --- |
|  |  |  | ***bla*_CTXM_** | ***bla*_OXA_** | ***bla*_CTXM 15_** | ***bla*_SHV_** | ***bla*_TEM_** | ***bla*_DHA_** | ***bla*_AmpC_** | ***qnrA*** | ***qnrB*** | ***qnrS*** | ***tetA*** | ***tetB*** | ***sul1*** | ***sul2*** | ***aac (3)-*IIA** |
| Cattle | Faeces | 92 | 61 (66.30) | 10 (10.87) | 7 (7.61) | 4 (4.35) | 51 (55.43) | 0 (0.00) | 89 (96.74) | 1 (1.09) | 8 (8.70) | 48 (52.17) | 10 (10.87) | 5 (5.43) | 2 (2.17) | 21 (22.83) | 6 (6.52) |
|  | Hand swabs | 6 | 5 (83.33) | 0 (0.00) | 0 (0.00) | 0 (0.00) | 4 (66.67) | 0 (0.00) | 6 (100.00) | 0 (0.00) | 0 (0.00) | 2 (33.33) | 0 (0.00) | 0 (0.00) | 1 (16.67) | 2 (33.33) | 0 (0.00) |
|  | Human stool | 3 | 3 (100.00) | 1 (33.33) | 0 (0.00) | 0 (0.00) | 1 (33.33) | 1 (33.33) | 0 (0.00) | 0 (0.00) | 0 (0.00) | 1 (33.33) | 0 (0.00) | 0 (0.00) | 1 (33.33) | 1 (33.33) | 0 (0.00) |
|  | Total | 101 | 69 (68.32) | 11 (10.89) | 7 (6.93) | 4 (3.96) | 56 (55.45) | 1 (0.99) | 95 (94.06) | 1 (0.99) | 8 (7.92) | 51 (50.50) | 10 (9.90) | 5 (4.95) | 4 (3.96) | 24 (23.76) | 6 (5.94) |
| Poultry | Faeces | 127 | 116 (91.34) | 15 (11.81) | 30 (23.62) | 1 (0.79) | 76 (59.84) | 0 (0.00) | 96 (75.59) | 0 (0.00) | 9 (7.09) | 36 (28.35) | 32 (25.20) | 9 (7.09) | 21 (16.54) | 40 (31.50) | 16 (12.60) |
|  | Hand swabs | 2 | 1 (50.00) | 0 (0.00) | 0 (0.00) | 0 (0.00) | 0 (0.00) | 0 (0.00) | 1 (50.00) | 0 (0.00) | 0 (0.00) | 0 (0.00) | 0 (0.00) | 0 (0.00) | 0 (0.00) | 0 (0.00) | 0 (0.00) |
|  | Human stool | 0 | NA | NA | NA | NA | NA | NA | NA | NA | NA | NA | NA | NA | NA | NA | NA |
|  | Total | 129 | 117 (90.70) | 15 (11.63) | 30 (23.26) | 1 (0.78) | 76 (58.91) | 0 (0.00) | 97 (75.19) | 0 (0.00) | 9 (6.98) | 36 (27.91) | 32 (24.81) | 9 (6.98) | 21 (16.28) | 40 (31.01) | 16 (12.40) |
| Grand Total | | 230 | 186 (80.87) | 26 (11.30) | 37 (16.09) | 5 (2.17) | 132 (57.39) | 1 (0.43) | 192 (83.48) | 1 (0.43) | 17 (7.39) | 87 (37.83) | 42 (18.26) | 14 (6.09) | 25 (10.87) | 64 (27.83) | 22 (9.57) |
